# Genome-wide development of miRNA-based SSR markers in *Cleistogenes songorica* with their transferability analysis to gramineae and non- gramineae species

**DOI:** 10.1101/723544

**Authors:** Gisele Kanzana, Yufei Zhang, Tiantian Ma, Wenxian Liu, Fan Wu, Qi Yan, Xueyang Min, Zhuanzhuan Yan, Blaise Pascal Muvunyi, Jie Li, Zhengshe Zhang, Yufeng Zhao, Jiyu Zhang

**Affiliations:** State Key Laboratory of Grassland Agro-Ecosystems; Key Laboratory of Grassland Livestock Industry Innovation, Ministry of Agriculture and Rural Affairs; Engineering Research Center of Grassland Industry, Ministry of Education; College of Pastoral Agriculture Science and Technology; Lanzhou University; Lanzhou 730020; People’s Republic of China

**Keywords:** *Cleistogenes songorica*, microRNA, simple sequence repeat, genetic diversity, transferability, target genes

## Abstract

SSR markers are commonly used for many genetic applications, such as map construction, fingerprinting and genetic diversity analysis due to their high reproducibility, levels of polymorphism and abundance. As endogenous, small RNAs, miRNAs have essential roles in plant development and gene expression under diverse stress conditions, including various biotic and abiotic stress conditions. In the present study, we predicted 110 pre-miRNAs sequences from 287 precursor miRNAs and used them as queries for SSR marker development. Among 110 primer pairs, 85 were successfully amplified and examined for transferability to other gramineae and non-gramineae species. The results showed that all 82 primer pairs yielded unambiguous and strong amplification, and across the 23 studied *Cleistogenes* accessions, a total of 385 alleles were polymorphic. The number of alleles produced per primer varied from 3 to 11, with an average of 4.69 per locus. The expected heterozygosity (He) ranged from 0.44 to 0.88, with an average of 0.74 per locus, and the PIC (Polymorphism Information Content) values ranged from 0.34 to 0.87, with an average of 0.69 per locus. In this study, 1422 miRNA target genes were predicted and analyzed using the GO (Gene Ontology) and KEGG (Kyoto Encyclopedia of Genes and Genomes) databases. The results showed that this miRNA-based microsatellite marker system can be very useful for genetic diversity and marker-assisted breeding studies.

---

*Cleistogenes songorica* (*C. songorica*), which belongs to the family Gramineae, is an important perennial forage and ecological grass in Northwest China, including in Inner Mongolia, where the average annual rainfall is 110 mm (Yang et al. 2001). It exhibits high feeding value, cold resistance and drought tolerance (Zhang et al. 2014) and has been domesticated as a turf grass cultivar *C. songorica* (Roshev.) cv. Tenggeli. To study the drought tolerance mechanism of *C. songorica*, leaf and root expression sequence tag (EST) resources have been used to investigate drought stress-responsive genes (Zhang et al. 2011). Some of these genes have been transferred into *Arabidopsis thaliana* and alfalfa to confirm and enhance stress tolerance in these plants (Duan et al. 2015; Zhang et al. 2016).

Small RNAs or microRNAs (miRNAs) are a class of noncoding RNAs 18-24 nucleotides in length. Endogenous small RNAs have essential roles in plant development, phase transition (Khraiwesh et al. 2012) and gene expression under diverse stress conditions, including biotic and abiotic stress conditions, and in different developmental stages of the life cycle (Ganie and Mondal 2015; Singh et al. 2017). Like small nucleolar RNAs (snoRNAs), short interfering RNAs (siRNAs) and piwi-interacting RNAs (piRNAs), miRNAs are not translated into proteins but are often involved in the regulation of gene expression (Lin et al. 2008; Neilson and Sharp 2008). miRNAs are highly conserved in both plants and animals and have been found in plants, green algae, viruses, fungi, and older lineages of animals (Bartel and Bartel 2003; Saini et al. 2008). In addition, some species-specific miRNAs exist, which regulate various developmental and biological processes (Fahlgren et al. 2010). Molecular markers or DNA markers have become efficient tools for identify polymorphisms among different genotypes or genes (Jiang 2015) and are increasingly used in plant molecular research. For example, DNA fingerprinting can be used to detect polymorphisms among individuals and has become a fundamental tool for crop improvement via plant breeding methods (Ahmad et al. 2010). DNA markers can be categorized in two types: non-polymerase chain reaction-based markers that identify restriction fragment length polymorphisms (RFLPs) and polymerase chain reaction-based markers. The latter type includes single nucleotide polymorphism (SNP) markers (Jin et al. 2003), intron length polymorphism (ILP) markers (Zhang et al. 2017) random amplified polymorphic DNA (RAPD) markers, amplified fragment length polymorphism (AFLP) markers (Bandelj et al. 2004) and simple sequence repeat (SSR) markers (Baraket et al. 2011; Zhang et al. 2012; Min et al. 2017). All of these DNA-based markers have been used in various genetic studies. The selection of the appropriate markers depends on the study objectives. The abundance, low cost, high polymorphism, heritability, multi-allelic nature, distribution, reproducibility, distribution throughout the genome, ease of use and generally codominant nature of SSR markers make them highly suitable for genetic diversity studies (Wiesner et al. 2001; Wassom et al. 2008; Cloutier et al. 2011; Smulders and De Klerk 2011; Kessuwan et al. 2016).

SSR markers, known as microsatellites, are one of the most variable types of short repetitive elements of 1-6 bases (Chen et al. 2009; Sujatha 2013) and are found in prokaryotic and all eukaryotic genomes (Schlötterer 2000). SSRs have many important biological functions, such as the regulation of chromatin organization; DNA metabolic processes, gene activity and RNA structure (Li et al. 2004; Li et al. 2002; Haasl and Payseur 2012). SSRS exhibit high polymorphism, abundance, and genetic diversity and tend to be codominant (Ni et al. 2002; Noli et al. 2008; Liu et al. 2010; Liu et al. 2016; Parveen et al. 2016). SSRs are widely used for genetic diversity analysis, germplasm identification, comparative genetics analysis, phylogenetic analysis, QTL analysis, linkage mapping and marker-assisted selection (Rakoczy-Trojanowska and Bolibok 2004; Saha et al. 2006; Cavagnaro et al. 2010; Liu et al. 2010; Mondal et al. 2015). With the identification of increasing numbers of SSRs, SSRs are widely used to overcome the restrictions associated with other types of markers for genome mapping, fingerprinting, and population genetics studies as well as in molecular breeding. Despite the advances in miRNA and SSR development, there is a need for the development of miRNA-associated markers, i.e., miRNA-SSRs, to study traits in *C. songorica* and other species. In the present study, we identified miRNA-SSRs in full genomic sequences of *C. songorica* pre-miRNAs. The present study describes the first genome-wide development of miRNA-SSR markers based on *C. songorica* and their transferability to other species.

## Methods and materials

### Plant material and DNA extraction

Plant materials contained gramineae and non- gramineae species. Gramineae species are *Cleistogenes* keng, wheat (*Triticum aestivum)*, ryegrass (*Lolium perenne*), rice (*Oryza sativa*) and maize (*Zea mays*), while non- gramineae are *Arabidopsis thaliana*, *Medicago truncatula*, alfalfa (*Medicago sativa*), yellow sweet clover (*Melilotus officinalis*), common vetch (*Vicia sativa*) and soybean (*Glycine max*). All these 11 species were used to examine the transferability of Cs-miRNA-SSR primers; and the 23 *Cleistogenes* accessions were used to analyze genetic diversity. In 23 *Cleistogenes* accessions one is cultivated type and the other 22 are wild types. The 22 *Cleistogenes* accessions were obtained from China (in Gansu, Inner Mongolia and Shandong provinces) and only one accession is from Mongolia **(Table S1 in File S1).**

The young leaves of the plants were collected separately and were bulked as a single sample and used for genomic DNA isolation. *Cleistogenes songorica* samples were frozen in nitrogen liquid and stored at −80°C until DNA extraction. Genomic DNA was extracted from the young leaf tissues using the sodium dodecyl sulfate (SDS) method (Shan et al. 2011). The extracted DNA was detected by agarose gel electrophoresis then the samples were diluted with ddH2O to 50 ng/μl and stored at −20 °C. DNA quality and quantity were checked by 2% agarose gel electrophoresis and spectrophotometric measurement using a NanoDrop ND 1000.

### Identification of SSRs, miRNA-SSR primer design *for C. songorica* and chromosome mapping

A total of 287 pre-miRNAs of *C. songorica* were extracted from the *Cleistogenes songorica* local genome and used for the identification and extraction of SSRs by using a Perl 5 script (*MISA*, MIcroSAtellite identification tool). Among the 287 pre-miRNAs, 110 pre-miRNAs sequences were selected and used as queries for designing primers flanking repeats. Flanking primers to the SSRs were designed using Primer3 software and Perl 5 interface modules. To design the SSRs in this study, pre-miRNA sequences with 100% matching (E value = 10^−10^) to the *C. songorica* genome sequence (local database in a webserver, www.biocloud.net) along with the 500 bp on each end (5’ and 3’) were used **(Figure S1 in File S2).**The minimum length criteria were 10 and six repeat units for mononucleotide and dinucleotide repeats, respectively, and five repeat units for trinucleotide, tetranucleotide, pentanucleotide and hexanucleotide repeats **(Table S2 in File S1)**. The miRNA-SSR primers were designed using BatchPrimer3, and the designed miRNA-SSR primers were synthesized by Shanghai Sangon Biological Engineering Technology (Shanghai, China). The primer design parameters were as follows: amplicon size, 100–350 bp; primer length, 18–27 bases with 20 as the optimum; annealing temperature, 57–63 °C with the optimum of 60 °C; GC content, 45%–50% (Liu et al. 2015). *C. Songorica* markers were mapped on 19 chromosomes using TBtools (https://github.com/CJ-Chen/TBtools) where 90 and 20 markers were successfully mapped and scaffolds respectively.

### Prediction of miRNA target genes and GO analysis

In the present study, we extracted pre-miRNAs from local genome database of *C. songorica* and after removing all the redundant, the corresponding pre-miRNA sequences of non-redundant pre-miRNAs were used as queries for a BlastN search using miRbase against the rice genome in grameneae (Jaiswal et al. 2002) to obtain the small mature miRNAs corresponding to the pre-miRNAs that were used as queries by using psRNATarget Server to identify miRNA target genes. Target genes were used for bioinformatics analysis, GO and KEGG pathway were analyzed using BioCloud software (https://www.biocloud.net).

### PCR amplification

PCR was carried out in a 10μl reaction mixtures containing 4.95 μl 2 × reaction Mix (dNTPs at 500 μM each, 20 mMTris-HCL, 100 mM MgCl_2_, 100 mM KCl_2,_ 3 mM MgCl_2_), 2.0 μl double distilled water, 1.0 μl Template DNA, 1.0 μl forward primers and 1.0 μl reverse primers, 0.05 μl Golden DNA polymerase. Amplifications were performed with pre-denaturation of 3min at 94 °C, 30sec at 94 °C, 30sec at 60 °C, 30sec at 72 °C, 30sec at 94 °C, 30sec at 56 °C and final elongation step of 7min at 72 °C. PCR products were visualized on 2% agarose gels using Gel Red staining or on 6% non-denaturing polyacrylamide gel using silver staining.

### Statistical analysis

The SSR marker profiles were scored in a binary format where alleles were indicated for absent (0) or present (1) of the corresponding bands among different *Cleistogenes* accessions. Individual bands that could be clearly scored were used for genetic diversity analysis. A dendrogram was constructed from a genetic identity matrix by using NTSYS-pc V.2.1 software and the UPGMA method. The significance of each node was evaluated by bootstrapping data over a locus for 1,000 replications of the original matrix. The genetic similarity analyses were performed using NTSYS-pc, and the pairwise similarities were obtained using Jaccard’s coefficients. The matrices of the resemblance coefficients were subjected to UPGMA to estimate the genetic similarity among the accessions and arrange the dendrogram. The genetic structures of twenty-three accessions were analyzed by using Cs-miRNA-SSR markers and STRUCTURE software. Principal components analysis (PCA) was used to compare the overall changes in the population structure of the accessions (Varshney et al. 2017). Ordination was performed with the ‘vegan’ package and plotted with the ‘ggplot2’ package of R statistical software. We transformed binary data from the amplified fragments of 23 accessions using Hellinger.

### Validation of PCR products by sequencing

To confirm the truly and exactitude of PCR amplification product and according to the presence of single bands and high amplification efficiency, we select some PCR amplification products and sequenced by the Shanghai Sangon Biotech Company. We used chromas software (http://technelysium.com.au/wp/chromas/) in order to obtain good and clear sequencing chromatograms nucleotide peaks results of PCR product sequences.

## Results

### The development and frequencies of miRNA-SSRs in the *C. songorica* genome

A total of 287 pre-miRNAs were predicted from the *C. songorica* local genome database, and 110 pre-miRNAs were used as queries to identify SSRs, yielding 125 SSRs. The MISA microsatellite search results revealed 0.88 kb distribution frequency of one SSR per locus. Examination of the SSR motifs in the SSRs containing pre-miRNA genes revealed that 27 pre-miRNA genes contained more than one SSR. Among the 125 total SSRs, 110 contained simple repeat motifs, whereas the remaining 15 contained compound motifs. Among the simple repeat motifs, mononucleotide motifs were most abundant (61, 48.80%), followed by dinucleotide motifs (46, 36.80%) and trinucleotide motifs (3, 2.40%) **(Table 1).** No tetra-, penta- or hexa-nucleotide repeats were found in any of the *C. songorica* pre-miRNA flanking sequences. In the 99 pre-miRNAs containing SSRs, mononucleotides occurred at the highest frequency (58%), being present in 58 miRNA genes, followed by dinucleotides (38%), present in 38 genes, and trinucleotides (3%), present in only 3 miRNA genes **(Figure 1A and B).**

**Table 1.** A summary of SSR search results.

| Search items | Numbers |
| --- | --- |
| Total number of pre-miRNA examined | 287 |
| Total number of identified SSRs | 125 |
| Number of SSR containing sequences | 99 |
| Number of pre-miRNA containing more than 1 SSR | 27 |
| Number of SSRs present in compound formation | 15 |
| Repeat type |  |
| Mononucleotide | 61 |
| Dinucleotide | 46 |
| Trinucleotide | 3 |
| Total length of sequences searched (kb) | 110 |
| Frequency of SSRs | One per 0.88kb |

**Figure 1.**
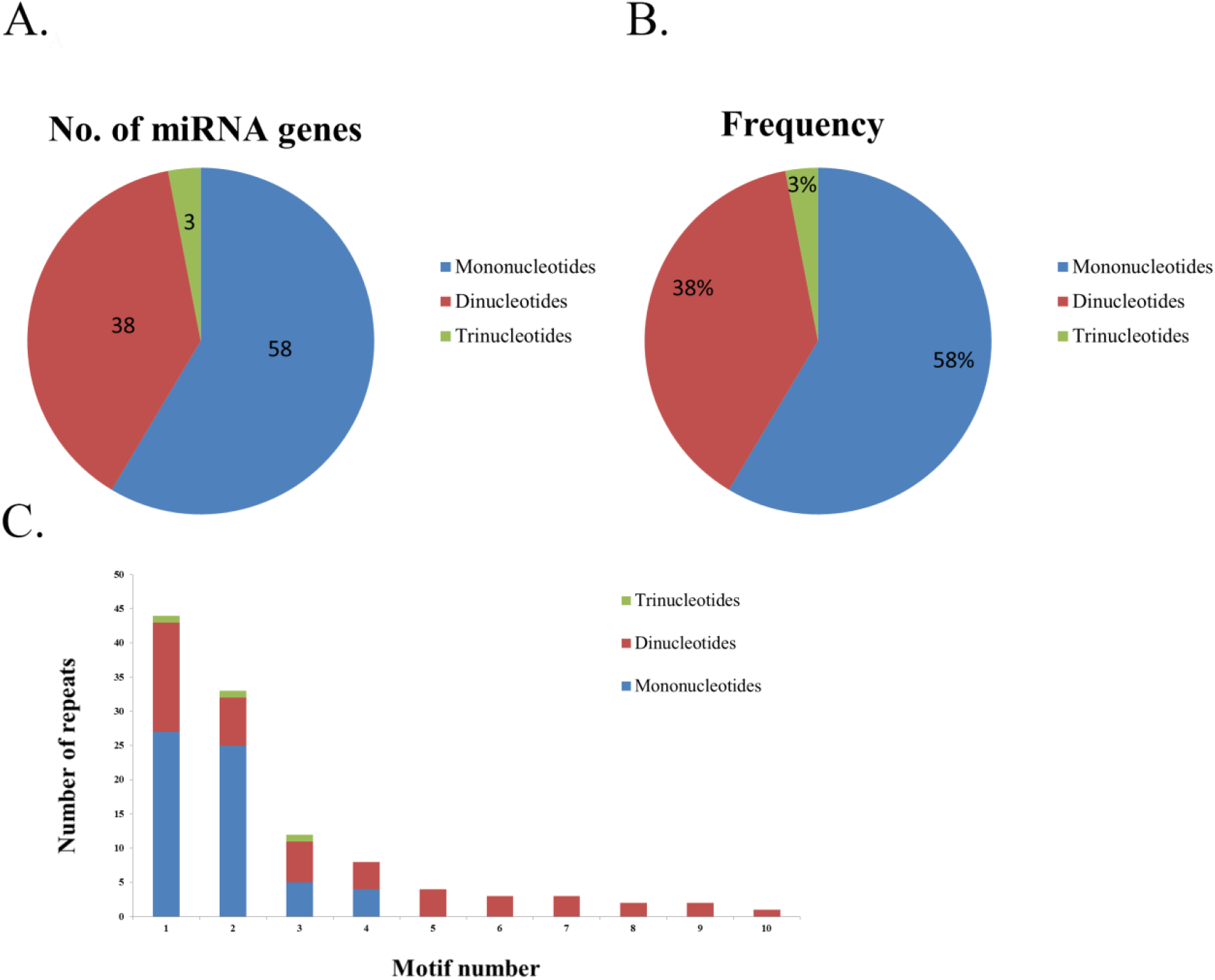
Number of miRNA genes with SSR motifs and abundance of miRNA-SSR motifs of *Cleistogenes songorica*. **A.** Number of miRNA genes possessing SSR motifs; **B.** Frequency of different repeat motifs in miRNA genes; **C.** Number of repeats having the most abundant miRNA-SSR motifs.

In the 110 SSR-containing miRNA genes, mononucleotide repeats were the most common type of repeat. Among the mononucleotides, (A)_27_ was occurred at the highest frequency (24.5%) in miRNA genes, followed by (T)_25_ (22.7%), (C)_5_ (4.5%) and (G)_4_ (3.6%). Among the dinucleotides, (CT)_16_ occurred at the highest frequency (14.5%) in the SSR-containing miRNA genes, followed by (TC)_7_ (6.3%), (AG)_6_ (5.4%), (TA)_4_ (3.6%), (GA)_4_ (3.6), (AT)_3_ (2.7%), (CA)_2_ (1.8%), (GT)_2_ (1.8%) and (TG)_1_ and (AC)_1_ (both 0.9%). Trinucleotide repeats occurred at low frequencies in the miRNA genes in the *C. songorica* genome, with (CCT)_1_, (TTG)_1_ and (AAT)_1_ each occurring at a frequency of 0.9% **(Figure 1C)**. Our study showed that mononucleotides were the most common type of repeat motif, whereas trinucleotides were the least common.

### Screening of Cs-miRNA-SSR primers, analysis of their transferability to other species and sequencing of PCR amplification products

All of the 110 Cs-miRNA-SSR primer pairs were screened for PCR amplification from the genomic DNA of *C. songorica*. Among the 110 primer pairs, 85 were successfully amplified, and 25 failed to be amplified. The 85 successfully amplified pairs were assessed for transferability to eleven gramineae or non-gramineae species. More than 70% of them yielded amplification products of the expected size (150-300 bp); the PCR products generated by the other primer pairs were smaller or larger than predicated. PCR products were then selected according to the presence of single bands and high amplification efficiency. For verification, PCR amplicons of several Cs-miRNA-SSR primers were amplified and sequenced. Among the sequenced alleles from the different miRNA-SSR primers, only some were homologous to the original locus from which the marker was developed. Primer pair Cs-miRNA-SSR-27, which contained trinucleotide repeats (TTG)_5_, and Cs-miRNA-SSR-82, which had dinucleotide repeats (TC)_11_, were matched to the locus used for marker development. Primer Cs-miRNA-SSR-108 and Cs-miRNA-SSR-67, which contained (A)_9_ and (T)_4_, respectively, were not homologous to the marker locus **(Figure S2 in File S2).** The highest amplification percentage (98%) was observed in *Cleistogenes songorica*, and the lowest (41.2%) was observed in *Glycine max*. The average amplification across the various species was 55.93% **(Table 2).** The amplification in *Arabidopsis thaliana* and *Vicia sativa*, which are non-gramineae species, was higher than in *Triticuma estivum* and *Zea mays*, two gramineae species. The 85 primer pairs were successfully amplified C. songorica were amplified in 11 species. C. songorica showed higher polymorphism than did the other species. A total of 466 alleles were detected by using 85 Cs-miRNA-SSR loci, ranging in frequency from 2 to 10 per locus. The three primer pairs numbered 33, 43 and 92 yielded the highest number of alleles (10), and the lowest numbers of alleles were obtained from ten primer pairs (nos. 7, 8, 11, 12, 17, 20, 23, 24, 28, and 36). The expected heterozygosity (He) ranged from 0.44 to 0.88, with an average of 0.74 per locus, and the PIC values ranged from 0.34 to 0.87, with an average of 0.69 per locus **(Table S3a in File S1).**

**Table 2.** Transferability of the 85 Cs-miRNA-SSR markers to the 11 gramineae and non-gramineae species

| No. | Genus/species | Transferability (%) |
| --- | --- | --- |
| 1 | *Cleistogenes songorica* | 98 |
| 2 | *Oryza sativa* | 64.7 |
| 3 | *Lolium perenne* | 60 |
| 4 | *Triticum aestivum* | 51.76 |
| 5 | *Zea mays* | 49.41 |
| 6 | *Arabidopsis thaliana* | 55.3 |
| 7 | *Vicia sativa* | 52.94 |
| 8 | *Medicago sativa* | 48.23 |
| 9 | *Medicago truncatula* | 44.7 |
| 10 | *Melilotus officinalis* | 47 |
| 11 | *Glycine max* | 41.2 |
| **Mean** |  | **55.93** |

### Genetic diversity, cluster and structure analyses of *Cleistogenes* accessions

The results showed that all 82 primer pairs yielded unambiguous and strong amplification, and across the 23 accessions, a total of 385 alleles were polymorphic. The number of alleles produced per primer varied from 3 to 11, with an average of 4.69 per locus. Cs-miRNA-SSR 45 and Cs-miRNA-SSR 75 had the lowest PIC values (both 0.41), and Cs-miRNA-SSR 43 had the highest PIC value of 0.86. The average PIC value was 0.62 **(Table S3b in File S1)**, which, being greater than 0.5, indicates the high level of polymorphism of these markers and suggests their potential for genetic diversity and genetic mapping analyses. To evaluate the application potential of the Cs-miRNA-SSR markers to the study of genetic diversity in *Cleistogenes songorica*, the 82 transferable primer pairs were analyzed in 23 *Cleistogenes* accessions. Due to the genetic closeness of *Cleistogenes songorica* with other *Cleistogenes* accessions, genetic marker array between *Cleistogenes songorica* and other *Cleistogenes* accessions could be evaluated and further use for its genetic diversity analysis. The banding patterns of 23 *Cleistogenes* accessions obtained with Cs-miRNA-SSR-31 and Cs-miRNA-SSR-82 markers are portrayed in **(Figure S3 in File S2).**

**Figure 2.**
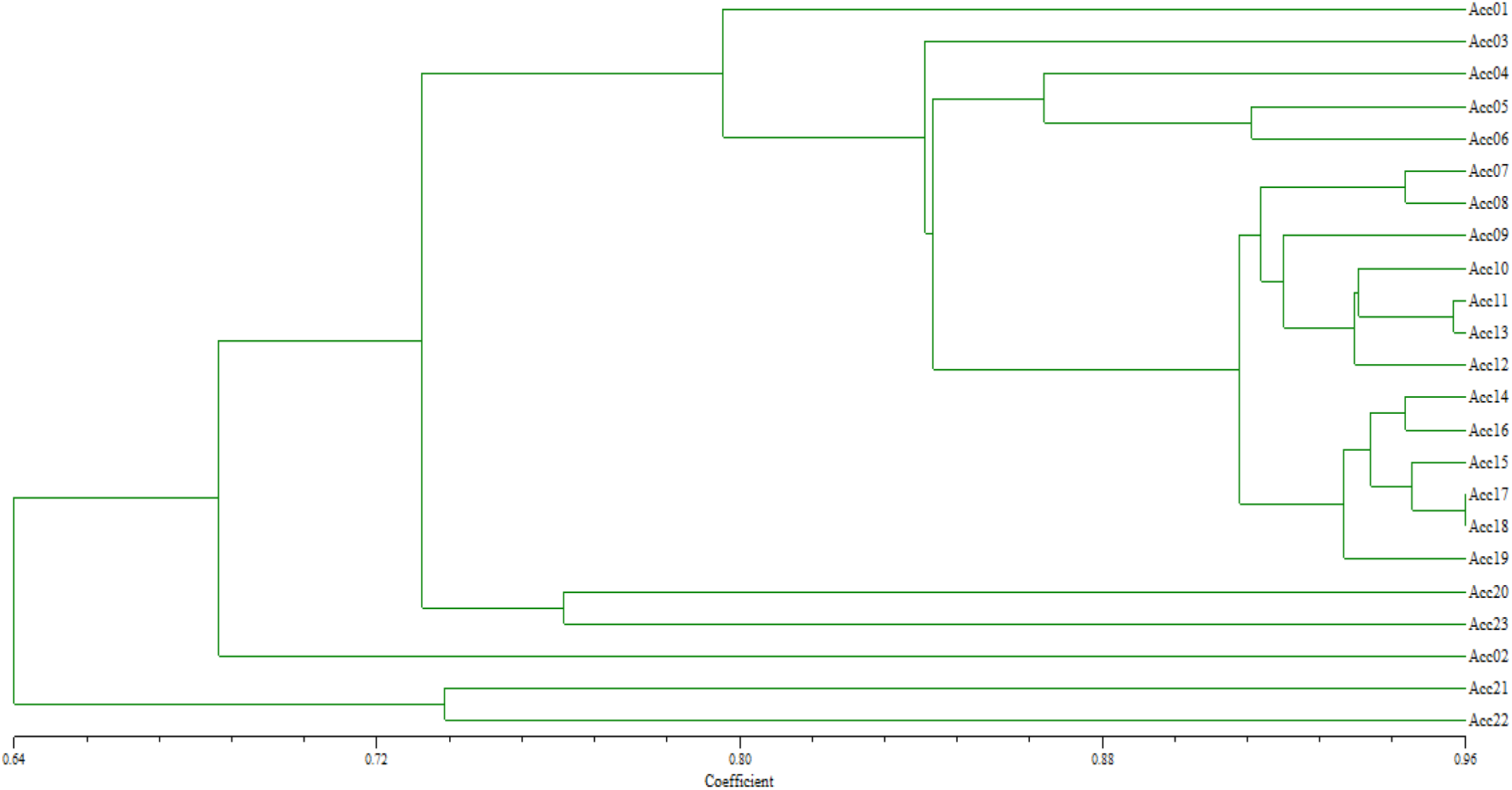
A dendrogram of unweighted pair group method arithmetic mean analysis (UPGMA) Analysis of the 23 *Cleistogenes* accessions grouped 17 of them together. In the cluster analysis, the twenty-three *Cleistogenes* accessions were divided into three clusters: cluster I, containing 20 accessions (Acc 01, Acc 03, Acc 04, Acc 05, Acc 06, Acc 07, Acc 08, Acc 09, Acc 10,Acc 11, Acc 12, Acc 13, Acc 14, Acc 15, Acc, 16, Acc 17, Acc 18, Acc 19, Acc 20 and Acc 23); cluster II, containing one accession (Acc 02); and cluster III, containing two accessions (Acc 21 and Acc 22) **(Figure 2).** In the analysis of population structure, which examined K=1-10, the optimal number of groups was three based on the maximum likelihood and delta K (∆K) values, consistent with the cluster results **(Figure 3)**.

**Figure 3.**
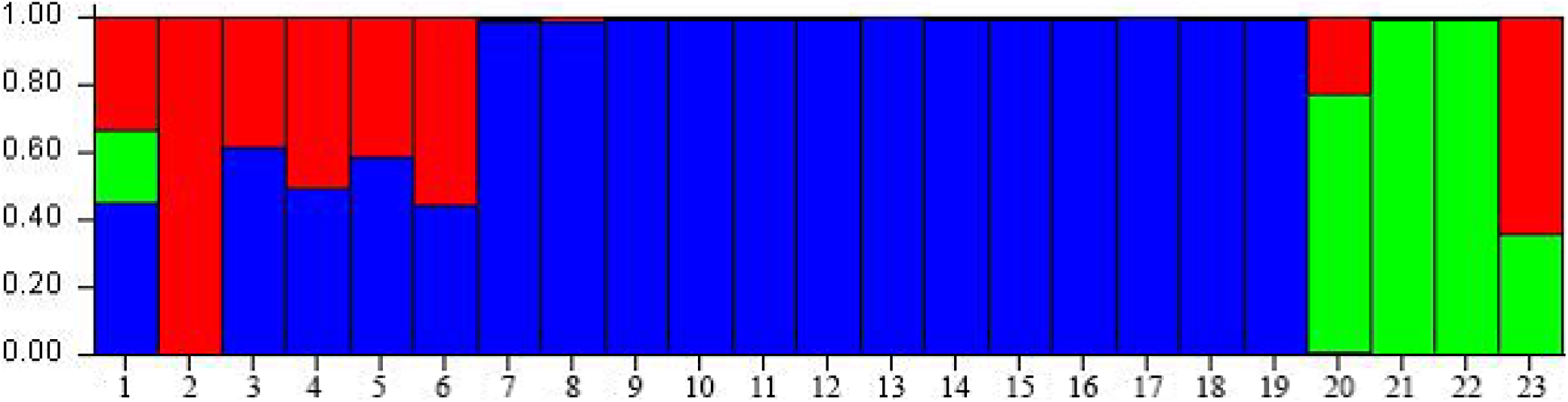
Histogram structure of the miRNA-SSR molecular markers data set for the model with K = 3 (showing the highest ∆K)

Among the three groups, group I contained eighteen accessions, group II contained one accession and group III contained 4 accessions. The PCA showed that 5 *Cleistogenes* accessions, including Acc 02 and Acc 20, which were *C. songorica*, did not cluster into any group, whereas the remaining 18 clustered together into one group. Most of these 18 were *C. songorica*, and one was *C. caespitosa* **(Figure 4)**. Although *C. hackelii* and *C. hancei* clustered in the same group, their genetic distance was large.

**Figure 4.**
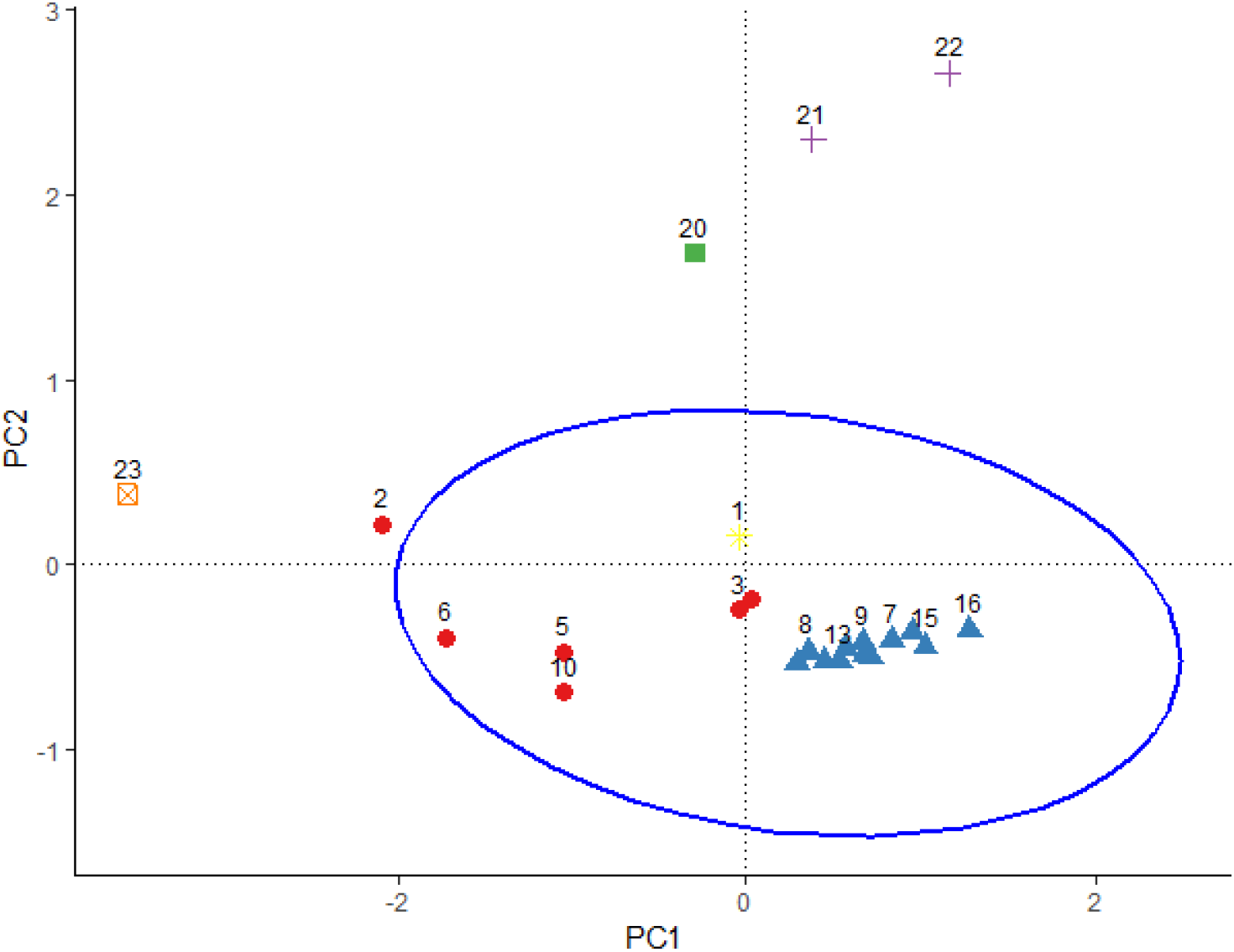
Grouping of geographical origin and genetic distributions of 23 *Cleistogenes* accessions using 82 pairs of miRNA-SSR markers. The principal components analysis (PCA) plot, each accession is represented by a vertical bar and the length of each colored segment in each vertical bar represents the proportion contributed by ancestral populations.

### Distribution of miRNA-SSRs on *Cleistogenes songorica* chromosomes

Among the 110 pre-miRNAs of the *C. songorica* genome, all 110 Cs-miRNA-SSRs were discovered. Although the 110 Cs-miRNA-SSRs markers were expected to physically map to 20 chromosomes, only 90 Cs-miRNA-SSRs were mapped, to 19 chromosomes. The remaining 20 Cs-miRNA-SSRs markers were located on scaffold regions, and no marker mapped to chromosome 3. Chromosome 12 contained the highest number of markers (12; 13.33%), whereas chromosomes 19 and 20 contained the lowest numbers of markers (1 each; 1.11%) **(Figure S4 in File S2).**

### Gene ontology and KEGG pathway analysis

To evaluate the potential functions of the miRNA target genes, GO and KEGG analyses were performed. Annotations of the 1422 predicted target genes were analyzed, and 1235 genes were annotated and classified based on GO terms. The target genes were classified into three categories: biological process (19 GO terms), cellular component (12 GO terms) and molecular function (11 GO terms) **(Figure 5).** In the biological process category, the two most overrepresented GO terms were metabolic process (648 genes) and cellular process (634 genes), followed by single-organism process (497 genes). In the cellular component category, the most overrepresented GO terms were cell part (786 genes) and cell (784 genes), followed by membrane (419 genes). In the molecular function category, the most overrepresented terms were catalytic activity (614 genes) and binding (600 genes).

**Figure 5.**
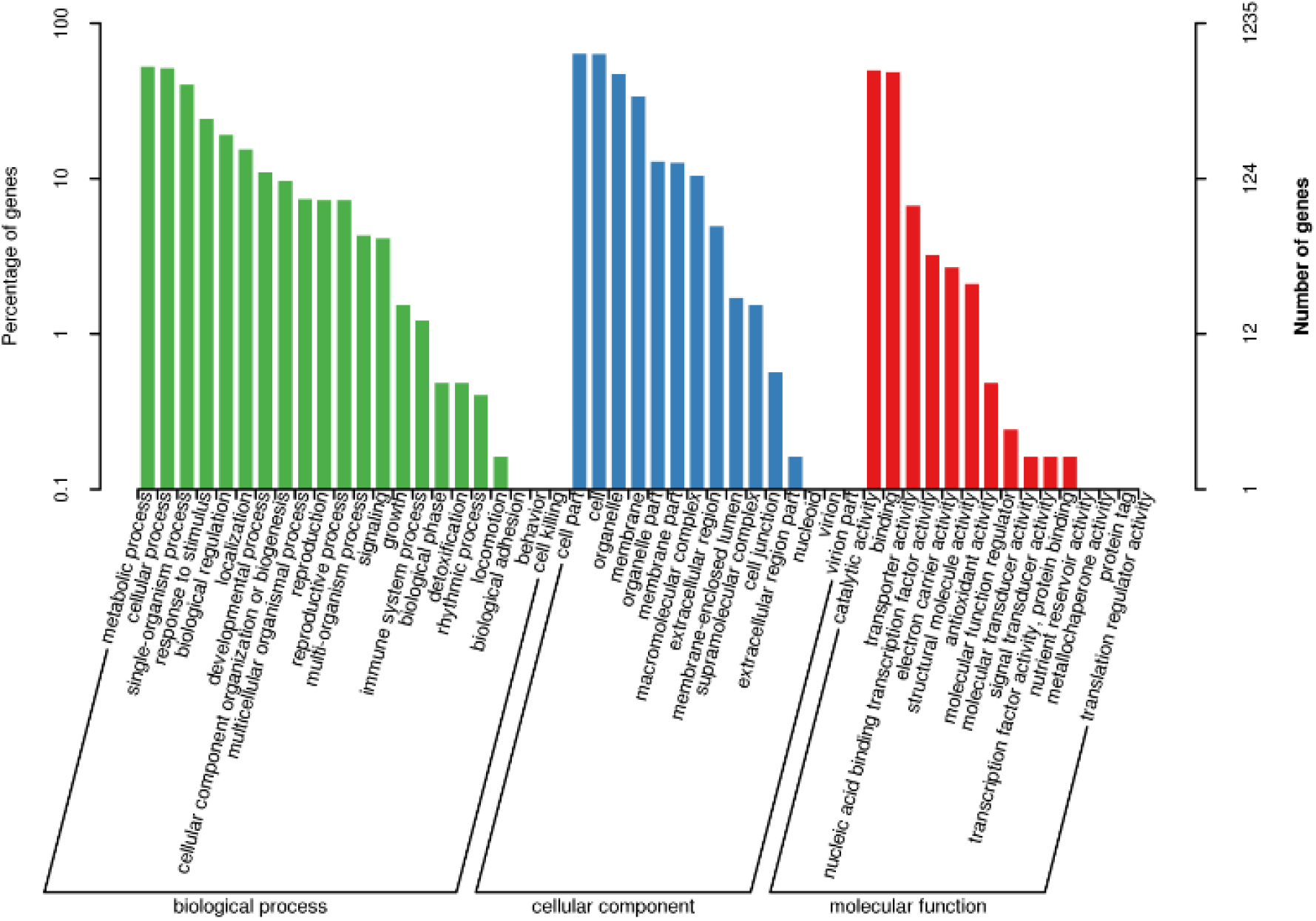
The relative frequencies of miRNA target genes assigned to the GO functional in 3 categories; biological process, cellular component and molecular function.

KEGG pathway analysis showed that 1422 miRNA target genes were annotated in the 20 most represented pathways. Enrichment analysis of the differentially expressed miRNAs identified five signaling pathways with q-values <1.00. Each enriched pathway is displayed in a scatter diagram, with points representing enrichment level, and the color corresponds to the number of genes enriched for the given pathway. A small *q*-value (red) indicates high significance of the pathway. Many genes were assigned to carbon metabolism, with others assigned to pyruvate metabolism, phagosome, glycolysis/gluconeogenesis and other pathways **(Figure S5 in File S2).**

## Discussion

In the present study, a total of 287 pre-miRNAs of *C. songorica* were predicted from the *Cleistogenes songorica* local genome. We discovered a total of 110 miRNA-SSRs from the 287 pre-miRNAs, and 110 pre-miRNA sequences were selected and used as queries for designing primers flanking repeats. In several studies, the flanking regions of pre-miRNA sequences, ranging from 300 bp to 1 kb, have been used to analyze various parameters of miRNA genes. To understand the origin and evolution of miRNA genes of ten different plants, including rice, (Nozawa et al. 2012) used the 300-bp flanking region at either end of pre-miRNA sequences. Similarly, to identify the single nucleotide polymorphisms in rice miRNA genes, (Liu et al. 2013) used the 1-kb flanking region from both ends of the pre-miRNA genes. Thus, we believe that the 500-bp regions on either side of the pre-miRNA sequences in this study form part of the miRNA genes that were used to design the SSRs in this study. Our study found no tetra-, penta- or hexa-nucleotides in the *C. songorica* pre-miRNA sequences. Among the repeat types, mononucleotide miRNA-SSRs occurred at the highest frequency, whereas trinucleotide miRNA-SSRs occurred at the lowest frequency. Among the 110 miRNA-SSR primer pairs, 85 were successfully amplified in 11 gramineae and non- gramineae species, and *C. songorica* showed the highest levels of polymorphism. The 82 transferable primer pairs were analyzed in 23 *Cleistogenes* accessions. All 82 primer pairs yielded unambiguous, high amplification and across the 23 accessions, and a total of 385 alleles were polymorphic. The number of alleles per primer varied from 3 to 11, with an average of 4.69 per locus. The highest PIC value was 0.86, and the average PIC value was 0.62. PIC values greater than 0.5 are considered to indicate informative markers, and loci with PIC values greater than 0.7 are suitable for genetic mapping (Bandelj et al. 2004). Of the 110 SSR-containing miRNA genes, mononucleotide repeats occurred at higher frequencies than did di- and tri-nucleotides. (A)_27_ mononucleotides occurred at the highest frequency (24.5%), and trinucleotides occurred at the lowest (all 0.9%). Our results indicate that the number of repeats is inversely related to repeat length such that as repeat length increases, number and relative percentage of SSRs decreases (Rakoczy-Trojanowska and Bolibok 2004; Ganie and Mondal 2015). Among the three groups, group I contained eighteen accessions, group II contained one accession and group III contained 4 accessions. The PCA showed that 5 *Cleistogenes* accessions, including Acc 02 and Acc 20, which were *C. songorica*, did not cluster into any group, whereas the remaining 18 clustered together into one group. Most of these 18 were *C. songorica*, and one was *C. caespitosa* **(Figure 4)**. Although *C. hackelii* and *C. hancei* clustered in the same group, their genetic distance was large.

The comparative electropherogram analysis of the four miRNA-SSR loci revealed that some of the sequenced alleles from the different miRNA-SSR primers were similar to the original locus from which the marker was developed, whereas others were not. Primer pair Cs-miRNA-SSR-27, with trinucleotide repeats (TTG)_5_, and Cs-miRNA-SSR-82, with dinucleotide repeats (TC)_11_, were matched to the locus used for marker development, but primers Cs-miRNA-SSR-108 and Cs-miRNA-SSR-67, which contained (A)_9_ and (T)_4_, respectively, were not. In previous studies, including those in *Melilotus* (Yan et al. 2017) and *Medicago truncatula* (Min et al. 2017), the sequenced alleles of SSR primers were similar to the original locus in cotton (Jena et al. 2012). The cluster analysis of the genetic relationships among the 23 *Cleistogenes* accessions, performed by UPGMA of the 82 polymorphic Cs-miRNA-SSR markers, clustered the 23 accessions into 3 clusters. There was no significant relationship between the pattern of clustering of members of cluster 1 and geographical location. This result may have been due to the small number of markers or accessions from each geographical location. The populations of the germplasm clustered together, and the genetic similarity coefficient ranged from 0.64 to 0.96, indicating close genetic relationships among the 23 *Cleistogenes* accessions (Wang et al. 2007). Similar results have been reported in other plant species (Verma and Rana 2011; Wang et al. 2013; Singh et al. 2014; Zhou et al. 2014).

It can be concluded that miRNA-SSR markers of *C. songorica* have great potential for studies of genetic diversity and for transferability to other Gramineae and non-Gramineae species. We hope that this study will be helpful for marker-assisted genetic improvement, for genotyping applications and in QTL analysis and molecular-assisted selection studies for plant breeders and other researchers.

## Acknowledgements

The author gratefully acknowledge the financial support from National Natural Science Foundation of China (31572453), Special Fund for Agro-scientific, Research in Public Interest (20120304205), Program for Changjiang Scholars and Innovative Research Team in University (IRT_17R50), Fundamental Research Funds for the Central Universities (lzujbky-2016-10), and the 111 Project (B12002).

